# Artemisitene protected against murine schistosomiasis japonica through anti-parasite activity and immune regulation

**DOI:** 10.1101/2023.08.11.552909

**Authors:** Meng-ke Liu, Xu-yang Chen, Juan-juan Tang, Zhi-peng Liu, Gui-ying Lin, Jun-ling Cai, Zuo-ming Chen, Yu-yun Yan, Xiao-fang Ji, Zhong-jin Yang, Zi Li

**Affiliations:** Sino-French Hoffmann Institute, School of Basic Medical Sciences, Guangzhou Medical University, Guangzhou 511436, China; School of Pharmaceutical Sciences and the Fifth Affiliated Hospital, Guangzhou Medical University, Guangzhou 511436, China; The Second Affiliated Hospital of Guangzhou Medical University, State Key Laboratory of Respiratory Disease, Guangdong Provincial Key Laboratory of Allergy & Clinical Immunology, Guangzhou Medical University, Guangzhou 510260, China

**Keywords:** Artemisinin analog, Artemisitene, *Schistosoma japonica*, liver fibrosis, tegument, immune regulation

## Abstract

*Schistosoma japonicum* (*Sj*) infection induced liver granulomatous inflammation and fibrosis. As an active artemisinin analog, the implication of artemisitene (ATT) in schistosomiasis were unclear. Herein, we found that ATT significantly reduced the count of total adult worms and eggs, and increased the count of single males, injured the tegument in the surface of *Sj* adult worms & gynecophoral canal of males. The transcription of 98 genes in females and 48 genes in males were significantly changed, and these genes were closely related to cellular anatomical entity through gene ontology analysis. So, ATT might possess anti-parasite activity. Meanwhile, ATT treatment significantly lowered the level of glutamic oxaloacetic transaminase (AST) and glutamic pyruvic transaminase (ALT) in sera, the size of mesenteric lymph node, and granuloma, the collagen area and α-SMA expression level in the liver. Liver transcriptome and multi-cytokines analysis indicated its immune regulation effect. Flow cytometry verified that the count of eosinophils in the liver were significantly increased, while the frequency of neutrophils, M1/M2 and Th1/Th2 index were significantly decreased. Therefore, we provided strong evidence that ATT has therapeutic potential through *Sj* clearance and anti-liver disease. Tegument development injury and immune regulation including type 2 immunity enhancement might be the mechanisms.

**Author summary:** Currently, there were still 290 million people worldwide who were infected by *Schistosoma*, and the treatment for schistosomiasis relies majorly on the use of a single drug-praziquantel. In this study, we described for the first time that artemisinin-derived artemisitene (ATT), chemically remarkably different from praziquantel, possessed the therapeutic effects on murine schistosomiasis japonica. ATT displayed both anti-*Schistsosoma japonicum* and anti-liver inflammation & liver fibrosis effect. Through RNA-seq and scanning electronic microscope of adult female & male worms from hepatoportal veins with or without ATT treatment, we found that the mechanisms of ATT’s anti-parasites could be through injuring tegument development and then interrupting adult worms’ especially adult female worms’ clearance by immune cells such as eosinophils. Moreover, through RNA-seq of liver total RNA, ELISA of multi-cytokines in liver lysates and flow cytometry analysis of liver single cells, we found that the anti-liver diseases’ efficacy of ATT was associated with immune regulation especially type 2 immunity enhancement. Therefore, ATT possessed the therapeutic potential against schistosomiasis japonica and further researches were necessary for its future clinical use.

## Introduction

Schistosomiasis is a helminthic disease caused by *Schistosomes* and infects more than 300 million people in 78 countries worldwide, and *Schistosoma japonicum* (*Sj*) is the main causative agent in Southeast Asia [(WHO 2018), https://www.who.int/news-room/fact-sheets/detail/schistosomiasis]. The pathogenic life cycle of *Sj* started with cercariae penetrating into skin. Then, schistosomula was formed in the skin and later in the lung, and adult male & female worms were developed and released high number of eggs after 5 weeks of infection. Highly immunogenic substances released by deposited eggs in the liver tissue or secreted by adult worms accommodating in the hepatoportal vein elicited host immune responses, including granulomatous inflammation and fibrotic reactions. 6 weeks (6w) of infection induced high severity of granuloma & liver fibrosis which manifested as acute schistosomiasis japonica [1–4].

Innate and adaptive immune cells, including T helper cells (Th), eosinophils, neutrophils, and macrophages, and cytokines & chemokine from these cells exerted specific effects on hepatic pathology during acute liver schitosomiasis japonica [1, 3]. Neutrophils were believed to play an important role in *Sj* granuloma formation and liver fibrosis [5, 6]. Interferon-γ (IFN-γ) in mice contributed to neutrophil recruitment and neutrophil cleared large pathogens by releasing neutrophil extracellular traps (NETs) during the granulomatous response [6, 7]. *Sj* blocked NETs formation in wild-type mice (*WT*) by upregulating interleukin-10 (IL-10) expression [6]. Eosinophils, as an parasites’ clearance cell population, were not necessary for granuloma formation or liver fibrosis in mice following *Schistosoma mansoni* (*Sm*) infection, however, they are a prominent granulomatous component and an important source of IL-4 to maintain the Th2 responses [8, 9]. Macrophages were part of the mononuclear phagocytic system and were implicated in tissue homoeostasis and various infections. There were two main subtypes, classically activated macrophage or M1 with pro-inflammatory action, and alternatively activated macrophage or M2 with anti-inflammatory activity. The M1/M2 paradigm emerged as similar as Th profiles, Th1 and Th2. Macrophages were the most abundant in the liver granulomas of mice with *Sm* infection[1]. During the acute phase of *Schistosoma* infection (approximately 5-7 weeks post-infection), immune responses were largely of Th1 type, associated with increased numbers of M1 macrophages that produce IL-12, IL-6, TNF-α and nitrogen oxide (NO) [5, 10]. 6w of *Sj* infection induced moderate Th2 response, which was lower than 7w of infection [1, 11]. Excessive Th1 response led to severe acute cachexia followed by death, while Th2 immunity acted as a double-edged sword: anti-inflammatory effects and suppresses Th1-mediated immunopathology, or driving liver damage or even liver fibrosis. Therefore, maintaining Th1/ Th2 balance was important to control schistosomiasis.

Praziquantel, as the only drug for schistosomiasis’ clinical treatment, was still insufficient for the disease control and elimination because of its less effective on juvenile worms [12]. There is an urgent need for novel drugs. Artemisinin (ART) derivatives-dihydroartemisinin and artesunate were more stable than artemisinin and possessed antimalarial activities [13, 14]. Dihydroartemisinin, arteether, artemether & artesunate, which contain a unique endoperoxide bridge, were reported as effective against juvenile *Schistosoma* worms in schitosomiasis experimental model animals or clinical trials [15–18]. Therefore, combination therapy might be a good option for schistosomiasis control and elimination [15, 17, 19, 20]. Besides, artemisinin and their analogs showed anti-inflammatory, antibacterial, anti-tubercular, antiviral and anticancer activities [21] [22]. Nonetheless, the artemisinin and its analogs had potential neurotoxicity in *in vivo* animal models [23]. So, to reduce neurotoxicity, new artemisinin derivatives with high efficiency should be required. Artemisitene (ATT), kept the endoperoxide moiety, and contained α, β-unsaturated carbonyl structure, which improved some biologic activities.

Artemisinin-type compounds also played immunomodulatory effect during innate and adaptive immune responses in cell types and context-dependent manner, and they possessed immunosuppressive activity toward inflammatory and autoimmune diseases [24, 25]. In this study, we will check the role of ATT on schistosomiasis japonica, including its effect on both parasite and host, and identify its implication in host’s immunity.

## Results

### ATT possessed anti-Schistosoma japonicum activity in mice

Artemisinin-type drugs for schistosomiasis treatment still needs to be adequately addressed. In mice with schistosomiasis japonica because of ATT treatment (Figure 1A), the count of total *Sj* adult worms in the portal vein and mesenteric veins was significantly lowered compared to non-treatment (Figure 1B, p=0.026), and the count of single males was significantly increased (Figure 1B, p<0.0001). Consistently, inside the liver section, the count of deposited eggs was significantly decreased (Figure 1C, p=0.036, and Figure 1D).

**Figure 1.**
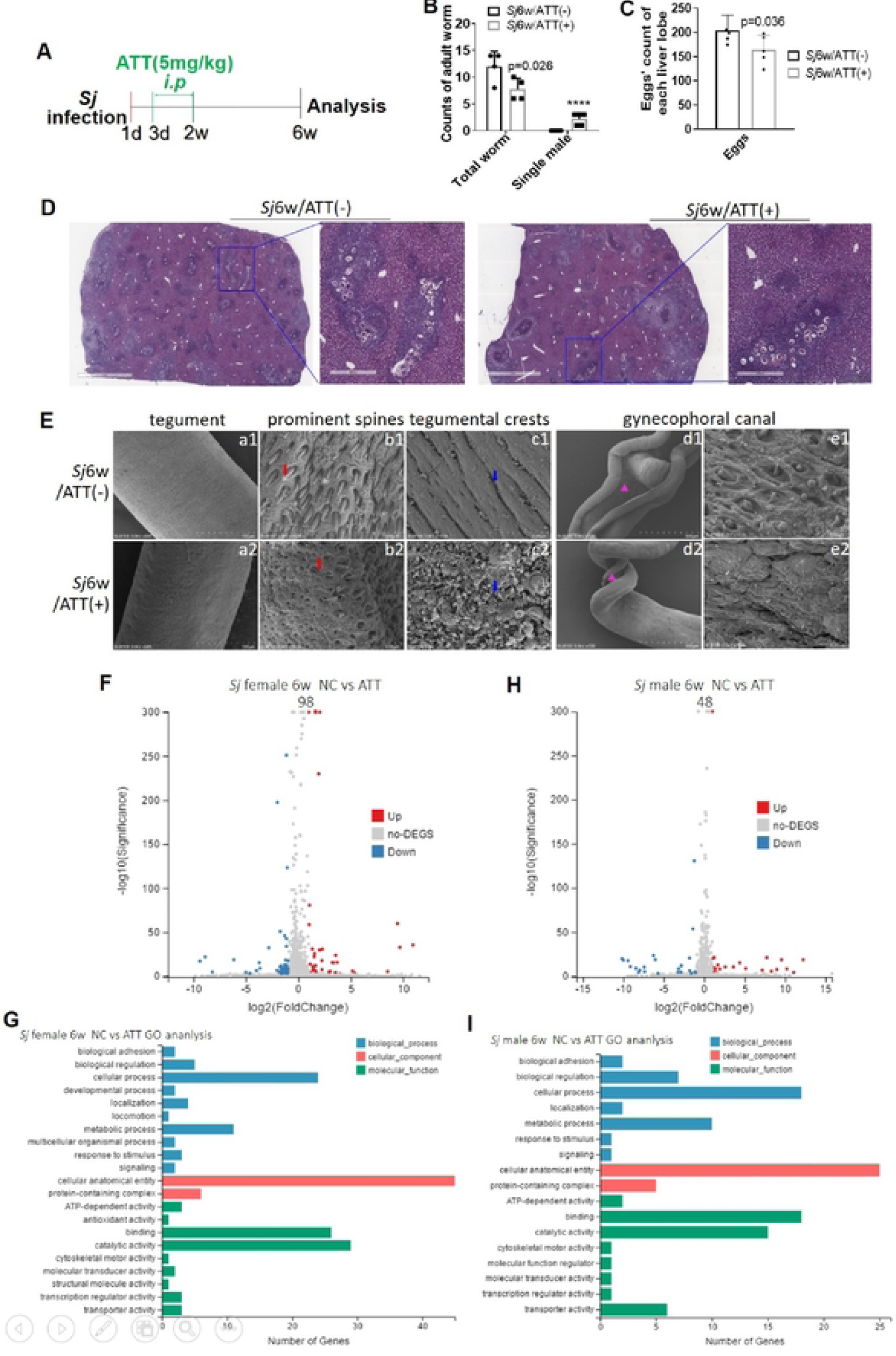
ATT possessed *anti-Schistosoma japonicum* activity in mice. **(A)** Flow chat of mice with *Sj* infection and ATT treatment. Mice were infected with *Sj* cercariae for 6 weeks, and ATT (5mg/kg body weight) was injected intraperitoneally on days 3-14 of *Sj* infection. (B) The number of total *Sj* adult worms and single male worms in the hepatoportal vein and mesenteric veins of mice with 6 weeks of *Sj* infection with or without ATT treatment was counted and analyzed (n=6, p=0.026). (C) The number of eggs in each liver section (2mm) of mice with 6 weeks of *Sj* infection with or without ATT treatment was counted and analyzed (n=5, p=0.036). And the typical liver sections (2mm or 400µm) of infected mice with or without ATT treatment after H&E staining were photographed and shown in (D). (E) Scanning electron microscopy (SEM) was used to show the surface of adult worms isolated from infected mice liver with or without ATT treatment. The tegument from the surface of adult worms’ mid-body with or without ATT treatment was shown in al or a2. The regular prominent spines (bl) and crests (cl) distributing in mid-body tegument, and prominent spines (el) in the surface of gynecophoral canal of adult males (d1) were injured by ATT treatment (b2-e2). (al & a2, 100µm; bl & b2, cl & c2 and el & e2, 5 µm; d1 & d2, 500µm). (F, G) Volcano plot of the differently expressed genes (DEGs) in *Sj* female adult worms (F) & male adult worms (G) from mice with or without ATT treatment was analyzed by using RNA-Seq analysis (FDR ≤ 0.001 and Log_2_ ≥ I). X-axis represents the log_2_ fold change and Y-axis represents the-log_10_(adjusted p-value). The threshold in the volcano plot was adjusted P value < 0.05 and |Log_2_ fold change| > I; Red and green represents up-regulated and down-regulated genes, respectively. Grey represents genes with no significant difference. (H, I) The significant Gene Ontology (GO) category for DEGs in *Sj* female adult worms (H) & male adult worms (I) from mice with or without ATT treatment. The X-axis and Y-axis represent −log_10_(p Valule) and the names of GO terms, respectively.

To further explain the anti-schistosomal efficacy of ATT in comparison with untreated control, adult female and male worms from mice was assessed by scanning electron microscopy (SEM) [26]. As shown in Figure 1E, both the mid-body surface of adult females and males, and the surface of gynecophoral canal of males showed various changes. Because of ATT treatment, apically directed spines were lost, the crests were ruptured. The tegument of gynecophoral canal exhibited severe damage, with the structure showing disorganization, collapse, and hole-shaped erosions. To investigate the differently expressed genes (DEGs) in adult worms after ATT treatment, worms were taken from the mice after 6w of Sj infection with or without ATT treatment for RNA sequencing (RNA-Seq). We obtained 20 million total sequence reads per sample. According to RNA-Seq analysis, the transcription of 98 genes in females (Figure 1F) and 48 genes in males (Figure 1H) were significantly changed by ATT treatment. Gene ontology (GO) enrichment analysis revealed that GO terms such as “cellular anatomical entity”, “catalytic activity”, “binding” and “cellular process” were significantly enriched in these changed genes, whereas most amount of genes were enriched into cellular anatomical entity (Figure 1G, 1I).

These revealed that ATT contributed to a reduction in female worms and then eggs number, and ATT directly damaged the surface structure of *Sj* & changing the transcriptional level of genes related to cellular anatomical entity might be the mechanisms.

### Artemisitene treatment reduced Sj infection-derived liver inflammation and fibrosis

Artemisinin and their analogs have shown anti-inflammatory and anti-fibrosis activity [22, 24, 27–31]. ATT decreased LPS-induced liver damage & inflammation through upregulating nuclear factor-erythroid 2 (NF-E2)-related factor 2 and its downstream protein heme oxygenase [32]. Mice with 6w of *Sj* infection manifested severe liver inflammation and fibrosis [4]. Herein, we found that 6w of *Sj* infection significantly increased the concentration of sera ALT and AST, and ATT treatment significantly decreased them (Figure 2A). *Sj* eggs-induced liver granulomas were classified into pre, early, mature and late stages according to H&E staining [1, 2]. ATT significantly lowered the area of granuloma induced by *Sj* eggs deposition, especially the area of early and mature stage granulomas (Figure 2B). We also found that ATT diminished the size of mesenteric lymph node (Figure 2C). These supported that ATT alleviated *Sj* infection-induced liver inflammation.

**Figure 2.**
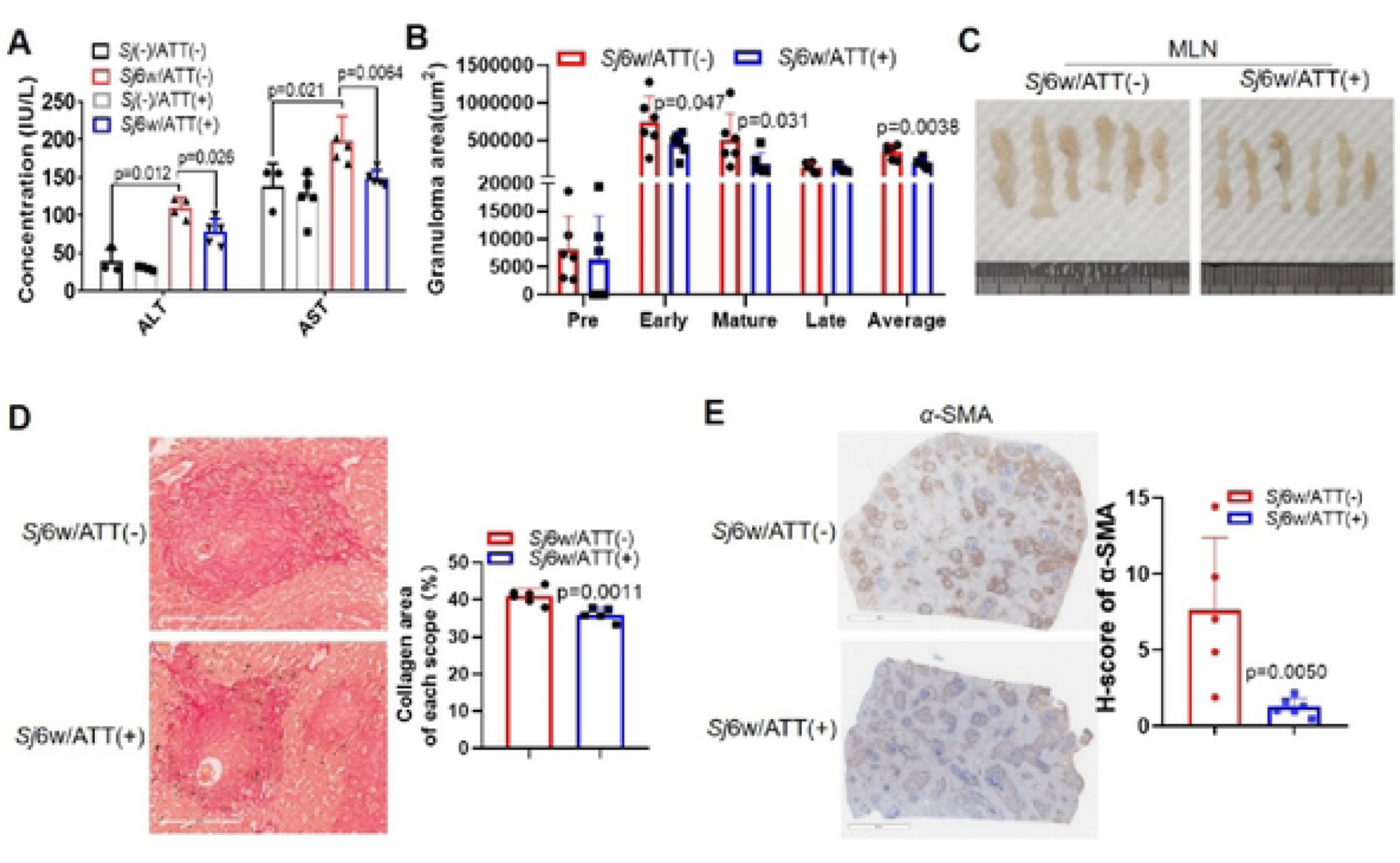
ATT significantly reduced *Sj* infection-derived murine liver inflammation and fibrosis. (A) The level of ALT and AST in the sera of indicated mice was tested and analyzed (n=4, without infection; n=5 & 6, with infection). (B) After H&E staining of liver sections, the percentage of area of granuloma, including pre early, mature and late stages was analyzed by Image J and then statistics. (C) Fresh mesenteric lymph node from indicated groups was photographed. **(D)** Typical Sirius red staining (200µm) of *Sj* single egg granuloma from liver of infected mice without or with ATT treatment was photographed (left) and percentage of morphometric collagen areas around *Sj* single egg (stained with strong red) was analyzed and shown (t-test, right). (E) The typical protein expression levels α.-SMA in mice liver tissue were determined by immunohistochemistry assay (IHC) and were shown (left), and the semi-quantitative analysis of this protein was analyzed using a modified H score (t-test, right). Data represent the mean± SD from different group experiments. Significant differences are indicated by *p < 0.05 and **p< 0.01.

According to Sirius red staining of the liver sections, ATT treatment significantly alleviated the collagen deposition area around single egg (Figure 2D, p=0.0011). In human, α-SMA marked activated HSCs and myofibroblasts in mice [33] [34]. The expression of α-SMA changed from moderate into low level according to H-score evaluation by IHC (Figure 2E). These supported that ATT treatment alleviated the extent of mice liver fibrosis because of 6w of *Sj*-infection.

### Both transcriptome and 23 cytokines’ assay of mice liver lysates indicated the immune modulatory function of ATT

To decipher the mechanisms of ATT’s effect on *Sj* infection induced liver inflammation & liver fibrosis, RNA-Seq and Bio-Plex Pro-Mouse Group I Cytokine 23-plex assay were used to screen the changes of transcriptome or some cytokines or chemokines in the liver lysates of mice with 6w of *Sj* infection with ATT treatment or not. RNA-Seq showed that 38 genes in the mice livers were significantly upregulated and 148 genes were significantly downregulated by ATT treatment when 6w of *Sj* infection (Figure 3A). Kyoto Encyclopedia of Gene and Genome (KEGG) pathway analysis of these changed genes revealed that most of these genes were enriched in immune modulatory activity such as antigen processing and presentation, viral protein interaction with cytokines, inflammatory bowel disease, NOD-like receptor signaling pathway, Th1 and Th2 cell differentiation, ect. (Figure 3B).

**Figure 3.**
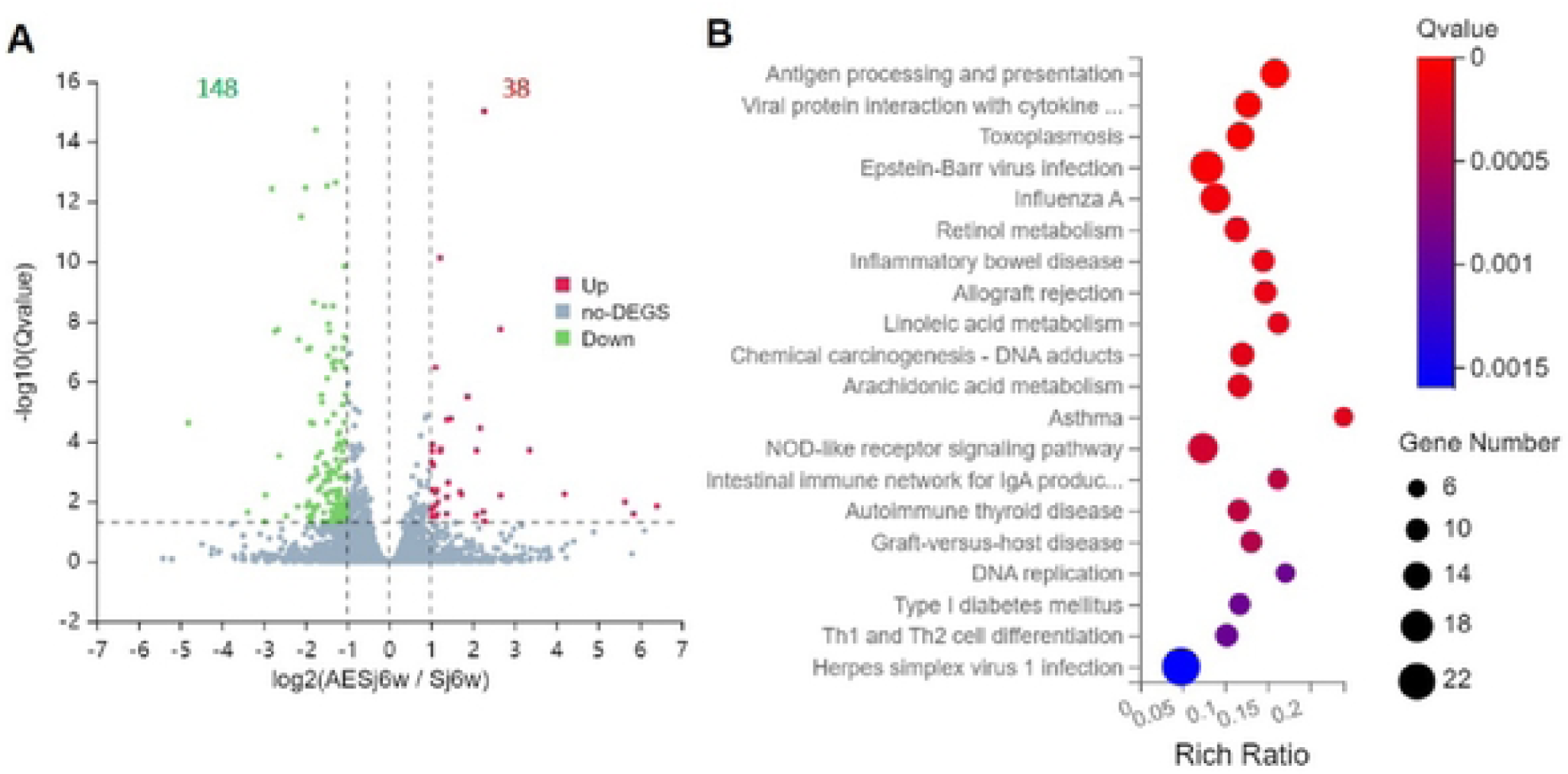
RNA-Seq suggested that ATT treatment affected innate and acquired immunity in the 6w of *Sj* infection mice liver. (A) Volcano plot of the DEGs of infected mice liver with or without ATT treatment. X-axis represents the log_2_ fold change and Y-axis represents the −log_10_ (adjusted p-value). The threshold in the volcano plot was adjusted P value < 0.05 and |Log_2_ fold change| > 1; Red and green represents up­regulated and down-regulated genes, respectively. Grey represents genes with no significant difference. (B) Kyoto Encyclopedia of Genes and Genomes (KEGG) pathway enrichment analysis showed the significant pathway for differentially expressed genes in infected mice liver with or without ATT treatment. The **X-axis** indicates −log_10_(p value) and the Y-axis represents the pathway name.

Bio-Plex Pro-Mouse Group I Cytokine 23-plex assay displayed that 6w of *Sj* infection significantly changed the level of 18 cytokines & chemokines. Proinflammatory cytokines in infected mice liver such as IL-1α, IL-1β, IL-6, TNF-α (Figure 4A), colony stimulative cytokines or chemokines such as G-CSF, KC, MCP-1, MIP-1α, MIP-1β, RANTES, Eotaxin (Figure 4B), activatory & differential cytokines (for innate lymphoid cells or T cell subsets) such as IL-12(p40), IFN-γ, IL-4 and IL-5 (Figure 4C) were significantly increased, while IL-3, GM-CSF and IL-17A were significantly decreased, while IL-2, IL-10, IL-13 and IL-9 did not show significant change (Figure4C). During ATT treatment the concentration of G-CSF, MIP-1α, MIP-1β, RANTES, Eotaxin (Figure 4B) and IL-12(p70), IL-4, IL-5, IL-10, and IL-9 (Figure 4C) in the livers of infected mice were significantly increased. Whereas the level of IL-1α, IL-1β, IL-6, TNF-α (Figure 4A); IL-3, GM-CSF, KC (Figure 4B); IL-12(p40), IFN-γ, IL-13 and IL-17A (Figure 4C) were not changed by ATT. These further supported the immune regulating effect of ATT such as type 2 immunity enhancement.

**Figure 4.**
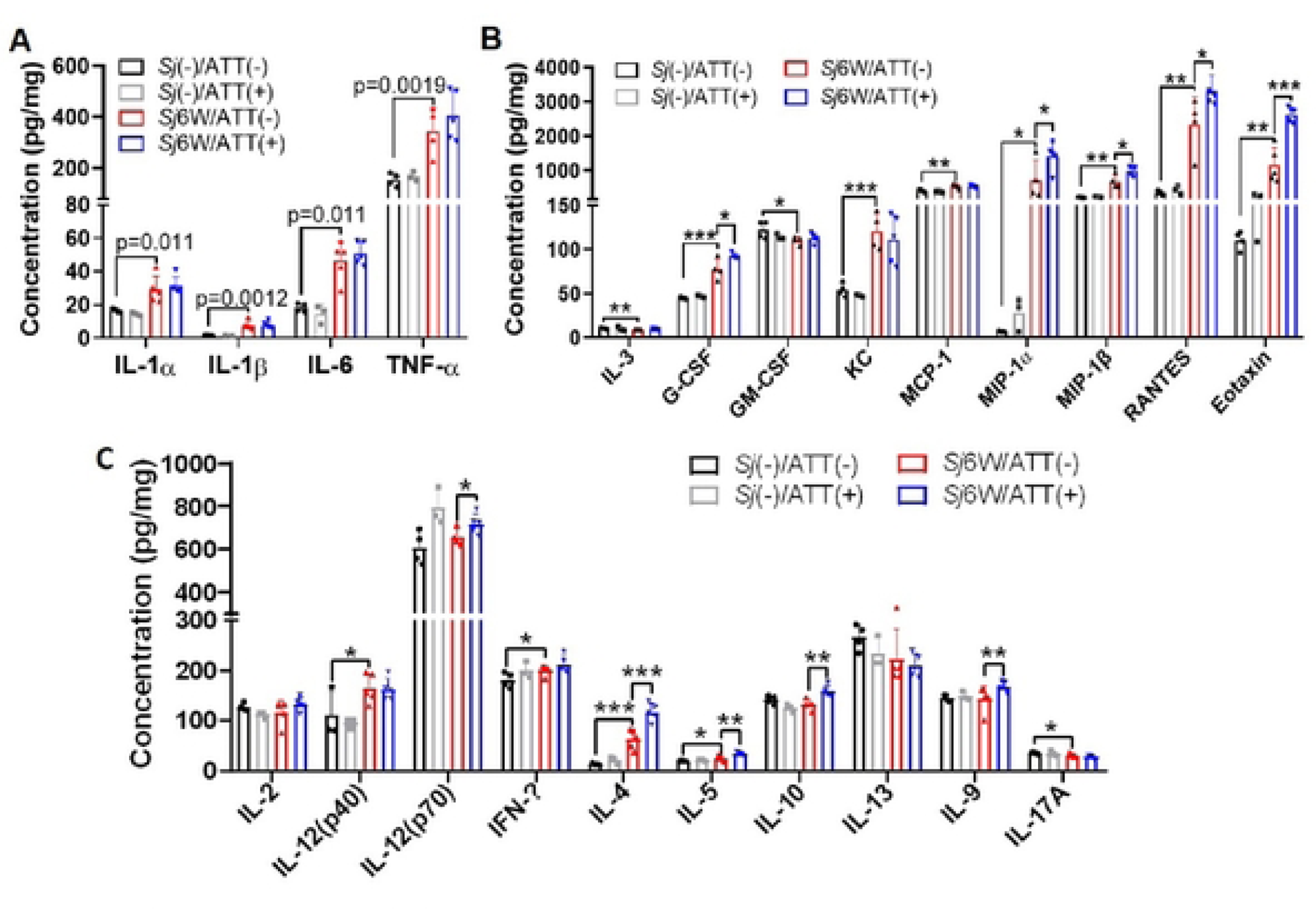
Concentration of immune molecules especially type 2 immune cytokines in infected mice liver was affected by ATT treatment. (A-C) The concentration of 23 cytokines including pro-inflammatorycytokines-IL-1α, IL-Iβ, IL-6, TNF-α (A), colony sti1nulative cytokines or chemokines-IL-3, G-CSF, GM-CSF, KC, MCP-1, MIP-1α, MIP-1β, RANTES, Eotaxin (B), and activatory & differential cytokines (for innate lymphoid cells or T cell subsets) such as IL-2, IL-12(p40), IL-12(p70), IFN-g, IL-4, IL-5, IL-10, IL-13, IL-9 and IL-17A (C) were tested by Bio-Plex Pro-Mouse Group I Cytokine 23-plex assay. Data represent the mean ± SD from different group experiments. Significant differences are indicated by *p < 0.05 and **p< 0.01.

### ATT decreased the percentage of neutrophils in the liver and spleen, M1/M2 index and Th1/Th2 index in the liver, and significantly increased the frequency of eosinophils in the liver and spleen of infected mice

Artemisinin analogs displayed an immunomodulatory effect during innate and adaptive immune responses in cell types and diseases context-dependent manner [25]. Neutrophils played an important role in *Sj* granuloma formation and liver fibrosis [5–7]. Eosinophils were a prominent *Sm* granulomatous component and an important source of IL-4 to maintain Th2 responses and they contributed to parasites clearance [1, 8, 9]. Macrophages were implicated in *Schistosoma* infections. Th1/ Th2 balance maintaining was important to control schistosomiasis, and M1/M2 paradigm emerged as similar as Th1/Th2 [1, 5, 10, 11]. To explore the effect of ATT on neutrophils, eosinophils, macrophages subpopulation, and Th1, Th2 subsets in *Sj*-infected mice liver or spleen at week 6 of *Sj* infection, mononuclear cells were isolated from mouse liver or spleen, and the percentage of CD45, CD11b and Ly6G co-expressed neutrophils (Figure 5A), CD45, CD11b and SiglecF co-expressed eosinophils (Figure 6A), CD45 and F4/80 co-expressed macrophages (MA) or its subpopulations with iNOS (M1) or CD206 (M2) expression (Figure 7A), and CD3^+^CD4^+^IFN-γ^+^Th1 & CD3^+^CD4^+^IL-4^+^Th2 (Figure 8A) were detected by flow cytometry (Figure 5-8).

**Figure 5.**
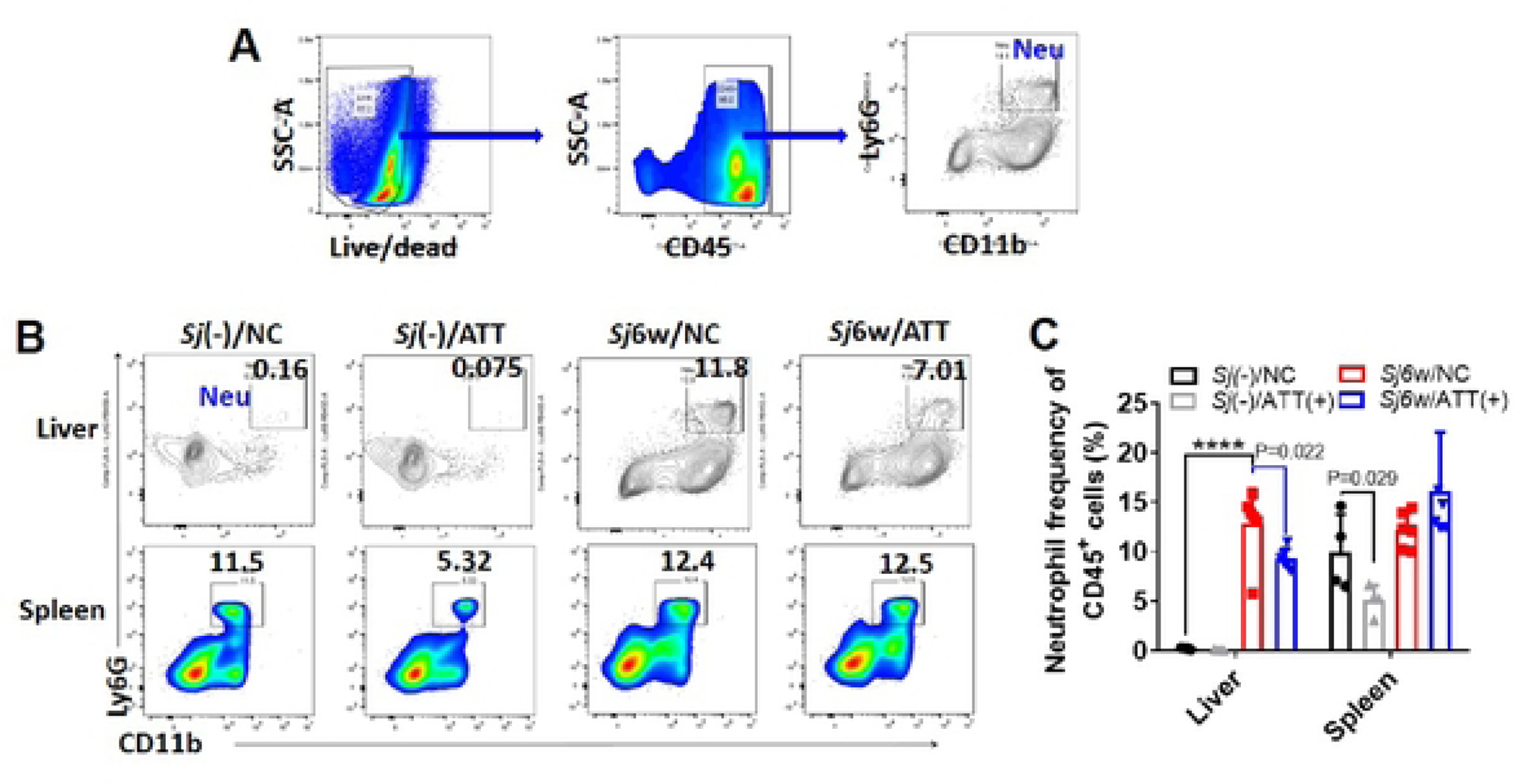
Neutrophils were significantly decreased in the liver but not in the spleen of infected mice with ATT treatment. Flow cytornetry was used for neutrophils analysis in the liver or spleen. Gate for nucleated cells was based on the size and complexity of the event (FCS-A and SSC-A, respectively). Nucleated cells were further plotted in FSC-A and FSC-H to gate single cells and exclude doublets. (A) From the single cell gate, live immune cells were defined as CD45+, and neutrophil gate was based on CD 11b^+^ Ly6G^+^. (B, C) The expression of neutrophils in CD45^+^ cells in the liver & spleen of infected mice with or without ATT treatment was evaluated. Typical neutrophils gating was shown in (B) and its quantification was shown in (C). Data represent the mean ± SD from different group experiments. Significant differences are indicated by *p < 0.05 and ****p< 0.0001.

**Figure 6.**
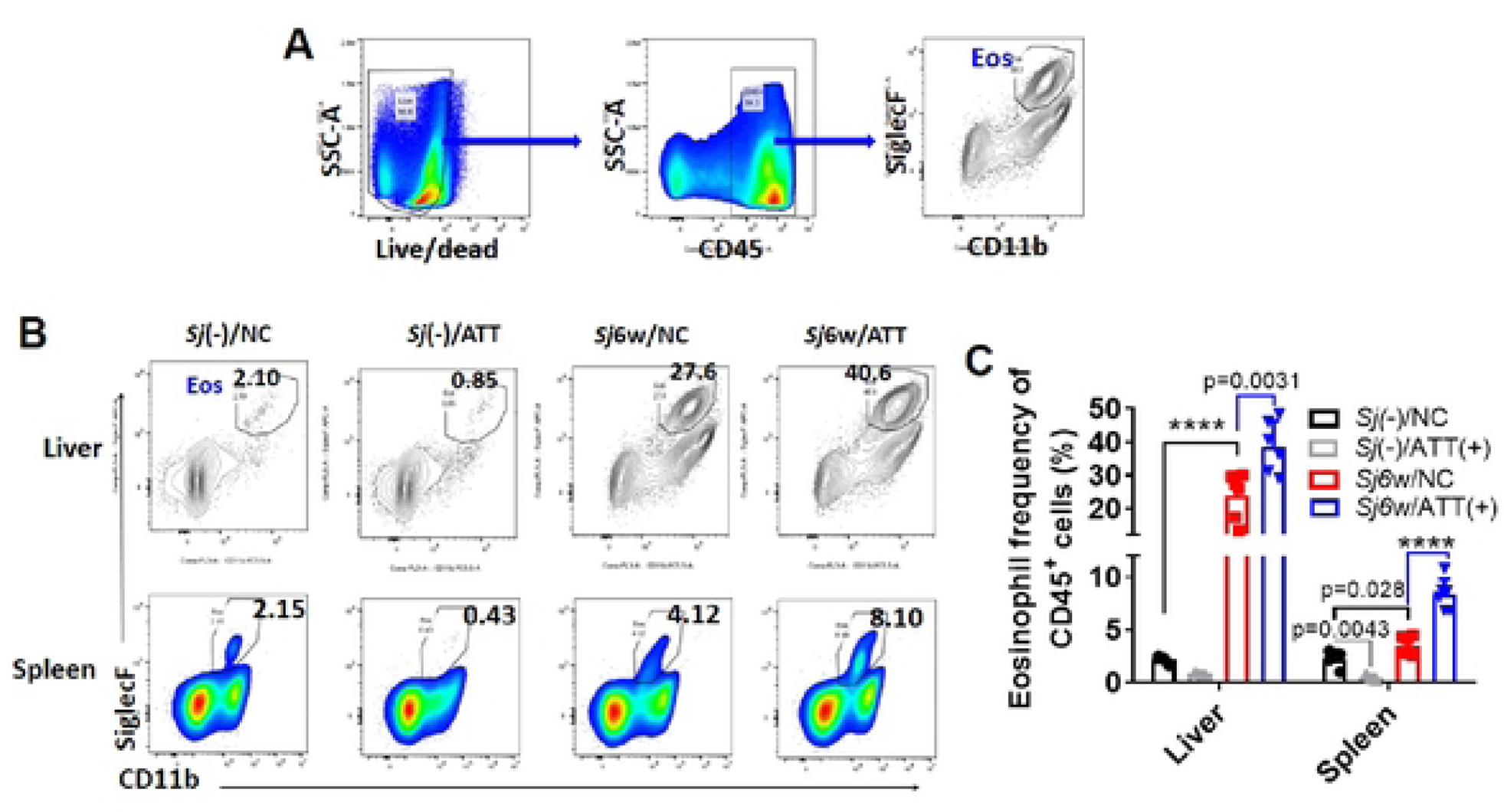
Eosinophils were significantly increased in the liver and spleen of infected mice with ATT treatment. The expression level of eosinophils in liver and spleen of mice was detected by flow cytometry. (A) From the single cell gate, gating strategy of eosinophil was CD45^+^CD11b^+^Siglecf^+^ in living cells. (B, C) The expression of eosinophils in CD45^+^ cells in the liver & spleen of infected mice with or without ATT treatment was evaluated. Typical eosinophils gating was shown in (B) and its quantification was shown in (C). Data represent the mean ± SD from different group experiments. Significant differences are indicated by *p < 0.05, **p< 0.01 and ****p< 0.0001.

**Figure 7.**
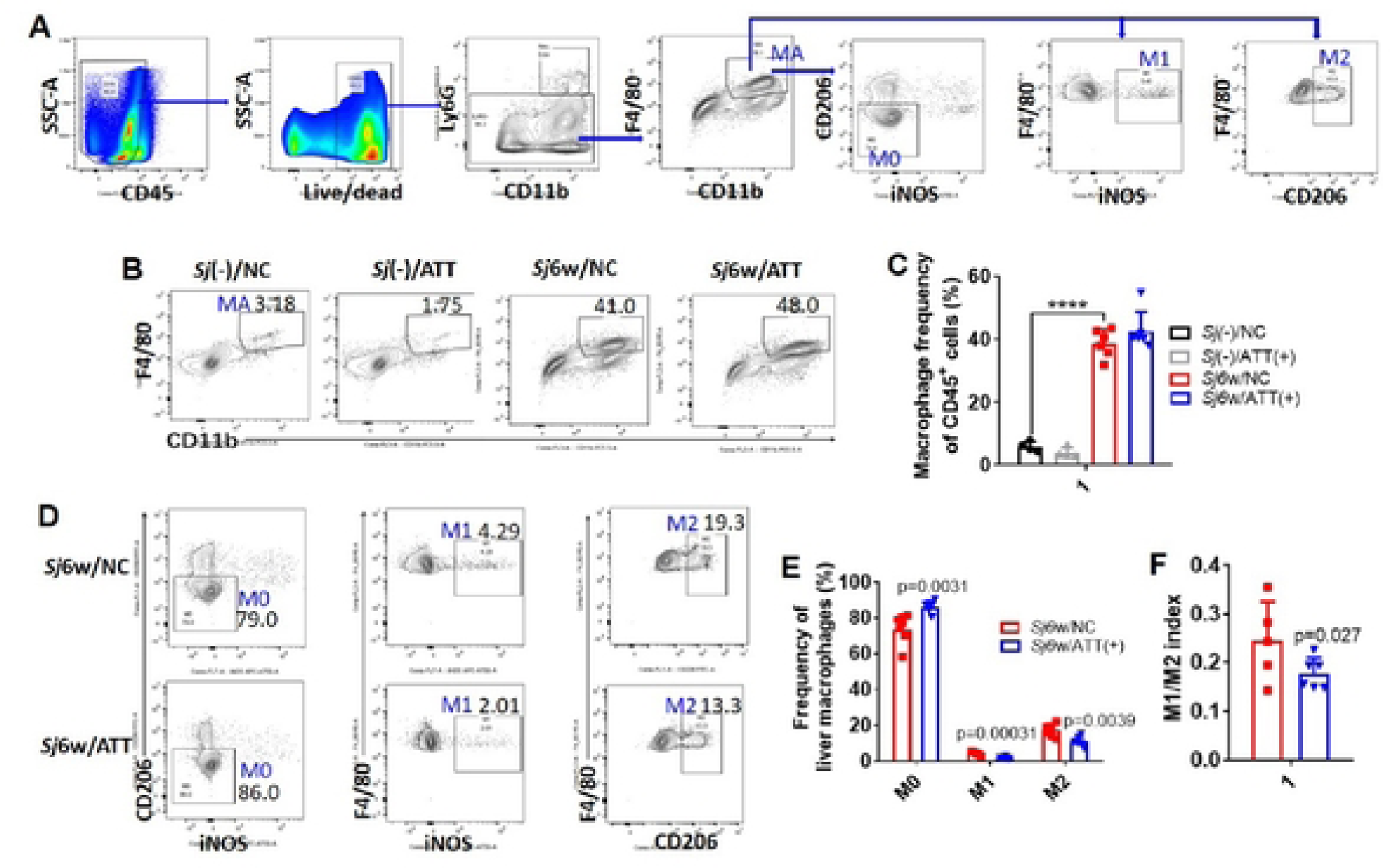
MJ/M2 index was significantly decreased in the liver of infected mice with ATT treatment. The expression levels of macrophages and its subpopulations in the liver of indicated mice were detected by flow cytometry. (A) Macrophage gating strategy was CD45^+^Ly6G^−^F4/80^+^ in living cells, and gated further according to CD206 and iNOS expression into M0 (CD206^−^iNOS^−^), M1 (iNOS^+^) and M2 (CD206^+^), respectively. (B, C, D, E) The expression of macrophages and its subpopulations in CD45^+^Ly6G· cells in the liver of infected mice with or without ATT treatment was evaluated. Typical macrophage or M0, M1 and M2 gatings were shown in (B) and (D) and there quantification was shown in (C) and (E), respectively. The ratio of M1/M2 (index) was shown in (F). Data represent the mean ± SD from different group experiments. Significant differences are indicated by *p < 0.05, **p< 0.01, and ***p< 0.001.

**Figure 8.**
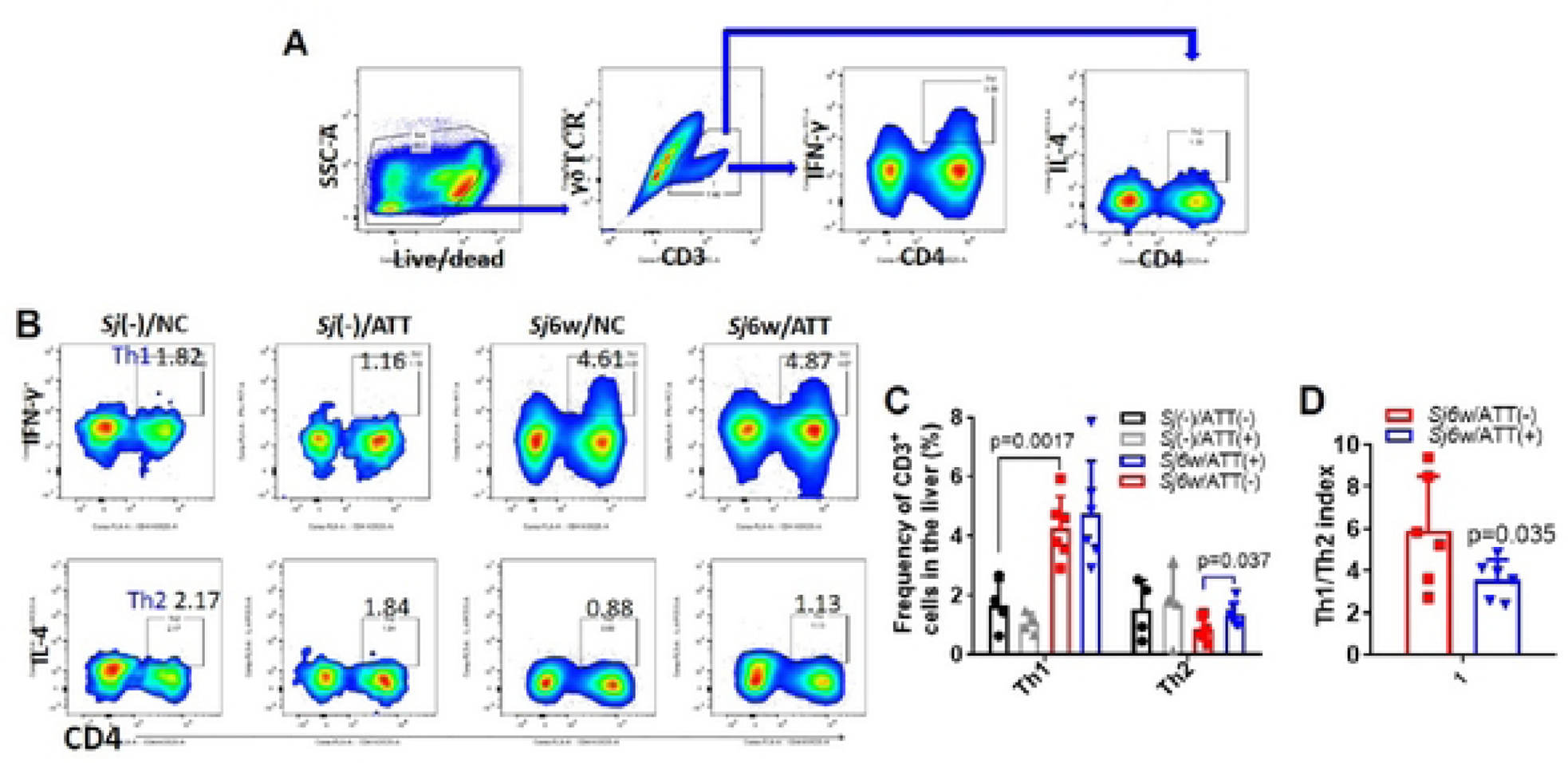
Th1/Th2 index was significantly decreased in the liver of infected mice with ATT treatment. (A) The expression levels of Th1 and Th2 in the liver of indicated mice were detected by flow cytometry, and the gating strategy was CD45^+^CD3^+^γδTCR^­^CD4^+^IFN-γ^+^ (Th1) /IL-4^+^ (Th2) in living cells. (B,C) The expression of Th1 (CD45^+^CD3^+^γδTCR^−^CD4^+^IFN-γ^+^) and Th2 (CD45^+^CD3^+^γδTCR^−^CD4^+^IL-4^+^) in the liver of infected mice with or without ATT treatment was evaluated. Typical Th1 and Th2 gatings were shown in (B) and there quantification was shown in (C). The ratio of Th1/Th2 (index) was shown in (D). Data represent the mean ± SD from different group experiments. Significant differences are indicated by *p < 0.05 and **p< 0.01.

**Figure 9.**
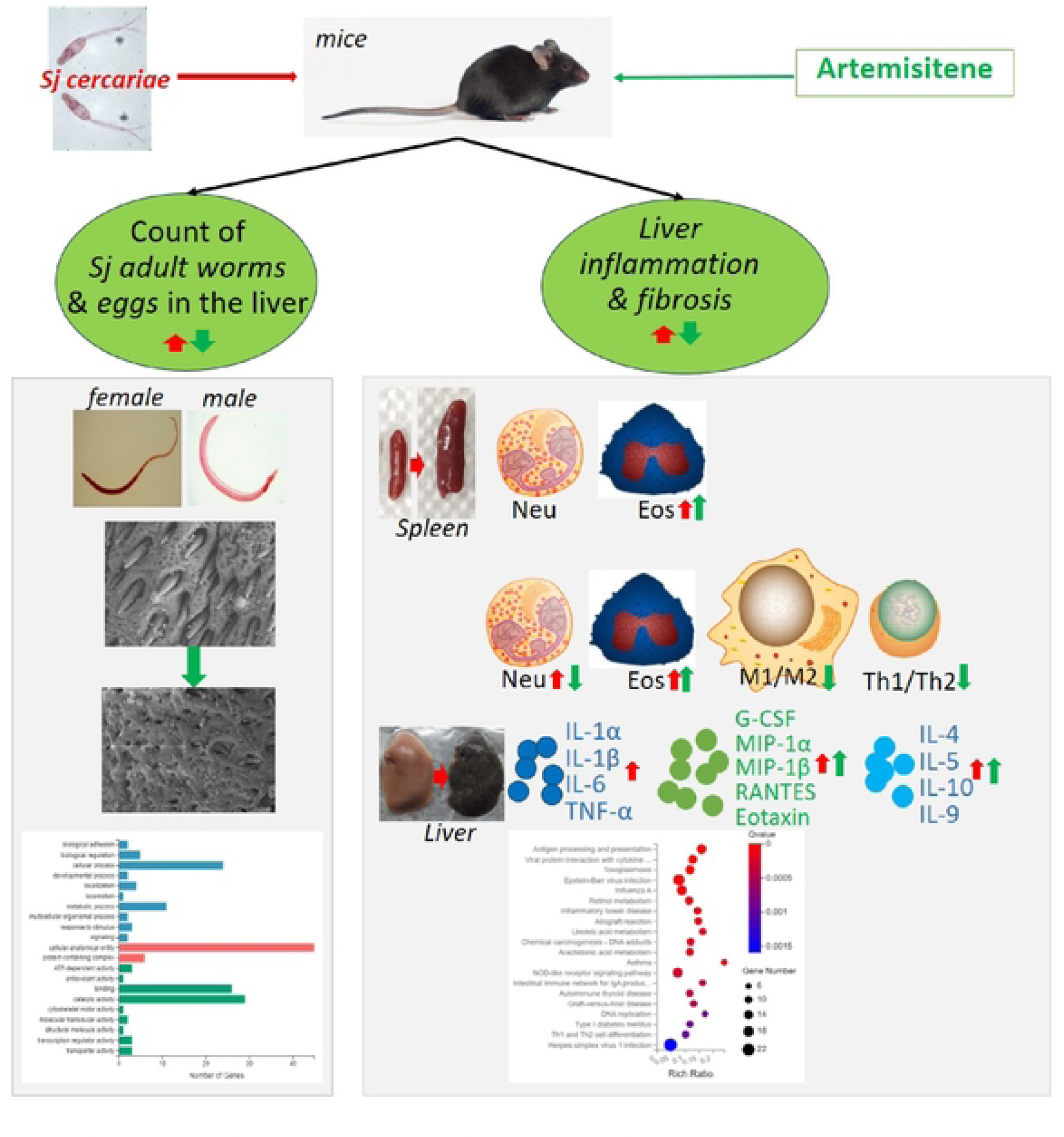
Graphical abstract & schematic model for the effect & mechanism of ATT on murine schistosomiasis japonica. During 6w of *Sj* cercariae infection, *Sj* adult female and male worms with normal tegument was developed and located majorly in hepatic portal vein and mesenteric vein, and liver granulomatous inflammation & fibrosis were induced because deposited *Sj* eggs and soluble antigen from adult worms & eggs stimulated neutrophils, eosinophils, macrophages and Th1 cells infiltration and production of some proinflammatory cytokines in infected 1nice liver such as IL-1α, IL-1β, IL-6, TNF*-*α, colony stimulative cytokines or chemokines such as G-CSF, MIP­1α, MIP-1β, RANTES, Eotaxin, and activatory & differential cytokines (for innate lymphoid cells or T cell subsets) such as IL-4, IL-5, IL-10, and IL-9. After ATT treatment, the count of *Sj* adult worms & eggs in mice liver, and the extent of liver inflammation & fibrosis were significantly reduced. Injuring tegument development, changing transcription of genes involved in cellular anatomical entity, & enhancing the immune clearance from the host (such as inducing eosinophils infiltrating into the liver & spleen) might serve as the mechanisms of ATT’s anti-*Sj* activity. Whereas reducing inflammatory neutrophils infiltration, M1/M2 & Th1/Th2 index, and lowering production of colony stimulative cytokines or chemokines such as G-CSF, MIP-1α, MIP-1β, RANTES, Eotaxin, and activatory & differential cytokines (for innate lymphoid cells or T cell subsets) such as IL-4, IL-5, IL-10, and IL-9 gave the clue of ATT’s immune regulation effect on the host and explained its role in liver inflammation & fibrosis.

Herein, our analysis showed that *Sj* infection significantly increased the percentage of neutrophil in the liver (Figure 5B, 5C, p<0.0001) and ATT treatment significantly decreased it (Figure 5B, 5C, p=0.022). However, the count of neutrophils in the spleen was not affected by *Sj* infection and then ATT treatment (Figure 5B, 5C), whereas ATT treatment significantly decreased the basal level of neutrophils in the spleen (Figure 5B, 5C, p=0.029). *Sj* infection also significantly increased the percentage of eosinophils in the liver (p<0.0001) and in the spleen (p=0.028) (Figure 6B, 6C), and ATT treatment significantly increase this cell population in the liver (p=0.0031) and in the spleen (p<0.0001) (Figure 6B, 6C), and ATT treatment also significantly decreased the basal level of eosinophils in the spleen (Figure 6B, 6C, p=0.0043). *Sj* infection significantly increased the amount of macrophages in the liver (p<0.0001), and ATT treatment didn’t affect the total amount of this cell population (Figure 7B, 7C). However, ATT treatment significantly increased the count of M0 (p=0.0031), and significantly decreased the count of M1 (p=0.00031) and M2 (p=0.0039) (Figure 7D, 7E), and M1/M2 index (Figure 7F, p=0.027). *Sj* infection significantly increased the amount of Th1 (p<0.0017) but not Th2 in the liver, and ATT treatment didn’t change the amount of Th1 but significantly induced Th2 (p=0.037) (Figure 8B, 8C). Th1/Th2 index was significantly reduced (Figure 8D, p=0.035), which was consistent to the effect on M1/M2 index (Figure 7F).

## Discussion

In this study, we firstly investigated the role and mechanisms of one of artemisinin analogs-ATT on murine schistosomiasis japonica. ATT not only showed anti-*Schistosoma* activity, but also displayed anti-inflammation and anti-fibrosis. Mechanically, ATT directly injured the surface of *Sj* and made the tegument of adult worm develop abnormally. Meanwhile, the tegument of gynecophoral canal of males exhibit severe damage because of ATT treatment, which might influence *Sj* male-female pairing and then eggs producing. Moreover, comparatively high eosinophils could exert the anti-parasite immunity. These explained the role and mechanisms of ATT’s anti-parasite effect. In terms of ATT’s anti-inflammation and anti-fibrosis activities, type II immunity enhancement might serve as the mechanism, since increased type 2 cytokines such as IL-4, IL-5 and IL-10, and decreased M1/M2 index & Th1/Th2 index will help tissue resist to *Sj* infection-induced liver inflammation & damage.

Some ART derivatives have shown the schistosomicidal potential [15–18]. ATT contained an extra structure-α, β-unsaturated carbonyl group compared to other ART analogs, which might strengthen this anti-parasite effect [35]. The anti-*Schistosoma* activity of ATT had not been previously tested. We firstly gave the solid evidence that early stage (3-14d post *Sj* infection) of ATT treatment manifested significantly anti-parasite and anti-liver disease effect on mice with 6 weeks of *Sj* infection. During ATT treatment, *Sj* was developing into *Schistosomulum* and locating inside the lung & in the bloodstream. Low dose of ATT treatment (5mg/kg body weigh) in early stage showed anti-worms and anti-eggs effect, which was not possessed by praziquantel. The lung, stomach and kidney of mice with ATT treatment didn’t display abnormal appearance (data not shown). Furthermore, when we treated mice with ATT during 4-6w of *Sj* infection, the count of *Sj* adult worms was also lowered (data not shown).

Morphologically, the teguments in the mid-body surface of adult worms or in the males’ gynecophoral canal by scanning electron microscopy were apparently injured by ATT treatment. According to RNA-Seq analysis, ATT treatment majorly changed the transcription of genes of male and female adult worms enriching in cellular anatomical entity. These gave the clue that ATT affected the tegument development. Whether ATT influenced other structures inside the parasites needed further study. Immune evasion effect has been found in the adult worms’ tegument [36]. The disruption of tegument will give more chance for eosinophils’ immunity against *Sj* in the murine liver.

Mice with 6w of *Sj* infection manifested as typical liver granulomatous inflammation and liver fibrosis, which has been widely used for therapeutic efficacy evaluation [1, 4] [2, 3, 37, 38]. The effect of ATT on liver disease and its immune regulation potential have not been reported. Herein, we found that ATT possessed anti-liver inflmammation and anti-liver fibrosis effect. Through RNA-Seq, multi-cytokines’ assay, and flow cytometry, we found that ATT enhanced type 2 immunity which might be the anti-disease efficacy mechanism. High eosinophil infiltration, type 2 cytokines including eotaxin, IL-4, IL-5 and IL-10 production, and low neutrophil infiltration, M1/M2 & Th1/Th2 index in the liver of mice with ATT treatment severed as the component of type 2 immunity. The immune regulating effect might be initiated by the changed development of *Sj* since ATT alone [*Sj*(-)/ATT(-) group] didn’t significantly change the basal level of cytokines and frequency of tested immune cells in the liver. However, ATT alone significantly decreased the basal frequency of neutrophils and eosinophils in the spleen. Although AST and ALT were significantly lowered by ATT, it didn’t significantly decrease the concentration of some proinflammatory cytokines such as IL-1α, IL-1β, IL-6 and TNF-α in the liver which were induced by *Sj* infection.

Taking together, we demonstrated with solid evidence that early stage of ATT treatment displayed anti-*Sj* activity though interrupting the tegument development in the surface of *Sj* & improving host’s anti-parasite immunity. Furthermore, ATT exhibited *in vivo* anti-liver inflammation and anti-liver fibrosis effect via enhancing type 2 immunity. Anyway, this study gave the clue to ATT’s therapeutic potential against schistosomiasis japonica, and further research was needed for its clinical trial in this infectious disease.

## Methods

### Reagents

ATT was produced and identified by professor Zhong-jin Yang’s group in the College of Pharmacy of Guangzhou Medical University[39]. Alanine transaminase (ALT) microplate assay kit (abs580004) and aspartate transaminase (AST) microplate assay kit (abs580002) were purchased from absin Biotechnology Co Ltd (Shanghai, China). Hematoxylin-eosin (HE) staining kit (BA-4025, Baso Biological Technology Co., Ltd, Zhuhai, China), Bio-Plex Pro-Mouse Group I Cytokine 23-plex assay (M60009RDPD, Bio-Rad, Hercules, California, USA), Sirius red staining kit was purchased from Abcam (ab150681, Cambridge, UK). Anti-alpha smooth muscle aorta (α-SMA) from Abcam (ab124964), anti-mouse-CD45-PE-Cyanine7 (30-F11, Cat no 25-0451-82), CD206-Alexia Flur^TM^ 488 (MR6F3, Cat no 53-2061-82), inducible isoform of nitric oxide synthase (iNOS)-APC-eFlur 780 (CXNFT, Cat no 47-5920-82), IL-4-PerCP-eFlur 710 (11B11, Cat no 46-7041-82), IFN-γ-PE-Cyanine 7 (XMG1.2, Cat no 25-7311-82), F4/80-PE, Dead-KO525, CD4-BV421, CD3-FITC and their corresponding isotype controls were obtained from eBioscience (San Diego, CA, USA). Anti-mouse-CD11b-PerCP-Cy^TM^5.5 (M1/70, Cat no 550993), SiglecF-Alexa Fluor 647 (E50-2440, Cat no 562680), Ly6G-BV421 (IA8, Cat no 562737) were from BD Biosciences (USA).

### Mice, parasite infection, and artemisitene treatment

6 to 8-week-old female C57BL/6 mice were obtained from specific pathogen free (SPF) Guangdong Medical Laboratory Animal Center and were maintained according to institutional guidelines. All mice experiments were approved as humane by the Institutional Animal Care and Use Committee at South China Agricultural University (2019-1013). Mice were infected by 20 ± 3 *Sj* cercariae of the Chinese main-land strain through abdominal skin penetration. ATT (5 mg/kg body weight) in 2% DMSO, 20% PEG300 and 78% saline was administered to each mouse by intraperitoneal injection once a day from day 3 to 14 of *Sj* infection (n=6), while the infection control group only received 2% DMSO, 20% PEG300 and 78% saline (n=6). Two non-infected control mice groups were treated with ATT (n=4) or 2% DMSO, 20% PEG300 and 78% saline (n=4), respectively. Mice were sacrificed at week 6 after *Sj* infection (Figure 1A). Sera, spleens, MLN, and liver tissues were collected for further analysis.

### H&E staining

Fresh hepatic tissues were fixed in 4% paraformaldehyde for 24 h and then were embedded with paraffin. Four-micrometer liver sections were prepared and stained with hematoxylin and eosin (H&E) to check the count of eggs, and assess granuloma size and the extent of liver granulomatous inflammation. The severity of liver granulomatous inflammation was evaluated according to the references [1, 40].

### Scanning electron microscopy of Sj adult worms

Ultrastructural features of the *Sj* female & male adult worms from mice with ATT treatment were examined using scanning electron microscopy (SEM) and were compared with those from untreated mice. For SEM, worms were washed three times in phosphate-bufered saline (PBS, pH 7.4) and fixed overnight at 4 °C in 2.5% glutaraldehyde-PBS solution (pH 7.4). Then *Sj* adult worms were washed again in PBS, and fixed in 1% osmium tetroxide (OsO4) for 1h, and dehydrated in an ascending series of ethanol. Finally, worms were dried, mounted on aluminum stubs, coated with gold, and examined using a Hitachi SU8100 SEM (Chiyodaku, Japan).

### Transcriptome analysis through RNA sequencing

Female or male *Sj* adult worms from mice portal vein & mesenteric vein, or liver tissues of mice with 6w of *Sj* infection with or without ATT treatment were harvested and washed twice with PBS. Total RNA was extracted and purified using TRIzol Reagent (Invitrogen) and then quantified and sequenced using the BGISEQ-500 platform by BGI Company (China).

### Immunohistochemistry staining

Endogenous peroxidase in sections from paraffin embedded mice livers was blocked with 3% hydrogen peroxide (H_2_O_2_). Immunohistochemistry staining (IHC) was used to determine the expression level and location of α-SMA in the mice liver tissue using anti-α-SMA (1:1000) primary antibody, followed by HRP-conjugated secondary anti-rabbit antibody. The images of 5 fields of every section were observed and captured with an optical microscope equipped with a camera (Olympus, Tokyo, Japan). The semi-quantitative analysis was determined by ImageJ software and shown as H-score [40, 41].

### Sirius red staining

Paraffin embedded liver tissues were cut into sections with 4µm thickness. These sections were used for Sirius red staining to evaluate the degree of liver fibrosis. The area of morphometric collagen as shown in red was analyzed using the Image J software. Each stained sample was evaluated in a double-blind fashion by two independent researchers.

### Multi-cytokines detection

Liver lysate samples were prepared using RIPA lysis buffer (P0013B, Beyotime Institute of Biotechnology, Shanghai, China) and stored at −80℃. Before analysis, they were left undisturbed at room temperature for 30 min. Cytokines were assayed through using Bio-Plex Pro-Mouse Group I Cytokine 23-plex assay kit (M60009RDPD, Bio-Rad, Hercules, California, USA) and the Bio-Plex® 200 System, MAGPIX Multiplex Reader (Bio-Rad Laboratories, Life Science Group 2000, Hercules, CA, USA). Briefly, the Luminex® xMAP® technology based on immunoassay methods is capable of simultaneously quantifying 23 targets: IL-1α, IL-1β, IL-2, IL-3, IL-4, IL-5, IL-6, KC, IL-9, IL-10, IL-12(p40), IL-12(p70), IL-13, IL-17A, Eotaxin (CCL11), granulocyte colony-stimulating factor (G-CSF), granulocyte-macrophage colonystimulating factor (GM-CSF), interferon gamma (IFN-γ), monocyte chemoattractant protein 1 (MCP-1), macrophage inflammatory protein-(MIP)-1α and MIP-1β, Regulated on Activation, Normal T Expressed and Secreted (RANTES, or CCL5), and tumor necrosis factor α (TNF-α). The concentration of each cytokine was extrapolated from the calibration curve (individual for each cytokine), determined independently for each experiment (each plate). The amount of samples from each group were more than 4.

### Preparation of single-cell suspensions of mice liver & spleen

Mice were anesthetized, and sterile normal saline was injected into the left ventricle to remove blood from organs. Then, the livers were cut and separated with forceps and digested in 3mg/mL Liberase (589994, Roche ^®^ Life Science) and 3mg/mL DNase I (Roche, 10104159001) for 1 hour at 37℃. The digested liver tissue pieces or cut spleen were rolled over a 100-um cell strainer (BD Falcon) to produce single cells and then suspending in Hanks’ balanced salt solution (HBSS). Red blood cells were lysed and removed with ammonium chloride for 10 min. Cell suspensions were incubated with LIVE/DEAD Zombie NIR ™ Fixable Viability Kit (Biolegend) for 20 min, and then re-suspended at 2–3×10^6^ cells/ml in complete RPMI 1640 medium with 10% fetal bovine serum (FBS).

### Cell surface & intracellular cytokines staining and then flow cytometry analysis

Cell suspensions were pre-blocked with mouse Fc block antibody (BD, clone 2.4G2). The following antibodies were used for cell surface marker staining: anti-CD45, CD11b, Ly6G, F4/80, SiglecF, CD206, iNOS; Or anti-CD3e, γδTCR, CD4. For intracellular staining of IL-4 or IFN-γ, cell suspensions were stimulated with phorbol 12-myristate 13-acetate (20 ng/ml; Sigma-Aldrich), ionomycin (1 μg/ml; Sigma-Aldrich), and BFA (10 μg/ml; Sigma-Aldrich) for 4 h at37 °C. Then, cells were fixed, permeabilized, and stained. Flow cytometry analysis was conducted in CytoFLEX (Beckman Coulter) and analyzed with FlowJo software (Ash-land, OR, USA).

Neutrophil was gated as CD45^+^CD11b^+^Ly6G^+^; Eosinophils: CD45^+^CD11b^+^SiglecF^+^; Macrophages: CD45^+^Ly6G^−^F4/80^+^, and macrophage subtypes such as M0: CD45^+^Ly6G^−^F4/80^+^iNOS^−^CD206^−^; M1: CD45^+^Ly6G^−^F4/80^+^ iNOS^+^; M2: CD45^+^Ly6G^−^F4/80^+^CD206^+^. T helper cell subtypes such as Th1 was gated as CD3^+^γδTCR^−^CD4^+^IFN-γ^+^ and Th2 as CD3^+^γδTCR^−^CD4^+^IL-4^+^.

### Statistical analysis

The results are presented as the standard deviation (± SD) of the indicated number of replicates/experiments. Data from each group were analyzed using SPSS software (v11.0) or Prism 8.0. Statistical evaluation of the difference between means was performed by one- or two-tailed, paired or unpaired Student’s t-test. A p-value of ≤ 0.05 was considered statistically significant.

## Acknowledgements

We would like to thank Prof. Shao-hong Chen and Yi Zhang from China center for disease control and prevention for supplying *Sj*-infected snails. This work was supported by the National Parasitic Resources Center, and the Ministry of Science and Technology fund (NPRC-2019-194-30), and the 111 Project (No. D18010).

## Author Contributions

ZL and ZY conceived the concept and critically reviewed the manuscript; ML and XC collected and assembled the data and finished the manuscript; ML, XC and GL performed the experiments; JC, ZL, ZC, JT, YY, and XJ provided experimental assistance. All authors contributed to the article and approved the submitted version.

